# Developing and Deploying a Scalable Computing Platform to Support MOOC Education in Clinical Data Science

**DOI:** 10.1101/2020.08.27.270009

**Authors:** David Mayer, Seth Russell, Melissa P. Wilson, Michael G. Kahn, Laura K. Wiley

## Abstract

One of the challenges of teaching applied data science courses is managing individual students’ local computing environment. This is especially challenging when teaching massively open online courses (MOOCs) where students come from across the globe and have a variety of access to and types of computing systems. There are additional challenges with using sensitive health information for clinical data science education. Here we describe the development and performance of a computing platform developed to support a series of MOOCs in clinical data science. This platform was designed to restrict and log all access to health datasets while also being scalable, accessible, secure, privacy preserving, and easy to access. Over the 19 months the platform has been live it has supported the computation of more than 2300 students from 101 countries.

## Introduction

One of the major challenges faced by data science educators is managing student computing environments. Typically educators must choose between teaching students how to set up an environment on their own computer or hosting a pre-configured server.^1^ Setting up a local environment on each student computer, while authentic, is challenging and often time-consuming because of the variety of operating systems and sometimes insufficient user permissions (e.g., for students using employer-provided computers). Server-based solutions shift managing the complexity of the computing environment to instructors, which reduces the authenticity of learning to manage the entire data science pipeline. A number of commercial solutions, like RStudio Cloud, have emerged to support educators providing a hosted solution without having to manage servers directly.^2^ While server-based solutions have some costs associated, for students with internet access they can increase equity of education as all students have equal computational power regardless of their own computing hardware.^1,3^

These technology challenges are magnified for those teaching data science focused Massively Open Online Courses (MOOCs). MOOCs are typically offered to thousands of learners across the globe completely asynchronously, increasing the number of unique computing environments and reducing instructor contact for individual-level support. Previous data science MOOCs have devoted an entire 4 week (∼13hour) course to setting up students’ computational environments.^4,5^ Others use technology embedded in the platform’s learning management system (e.g., shared JuptyerHub).^3,6^ Importantly, these data science MOOCs have not had a particular domain focus and thus can use openly available or non-sensitive data in their courses.

While MOOCs are an attractive solution to the increasing demand for a clinical data science workforce,^7^ it is not clear how to support student computing environments when working with sensitive healthcare data. We developed a series of MOOCs (“Specialization”) on clinical data science,^8^ that uses real clinical data (MIMIC-III demo database).^9^ At the time, all individuals seeking access were required to complete data use agreements. Setting up a student’s local environment would not allow us to restrict or track data download and potential sharing with external entities. Built in data science solutions on the course hosting site had the same limitation and additionally would not allow instructors to restrict data access to only those students who had completed a data use agreement. In response to these challenges we sought to create a hosted computing platform that would both manage student access to restricted materials and accommodate the unique challenges posed by MOOCs.

## Methods

### Designing the Computing Platform

The primary goal of developing the computing platform was to create a system that would allow instructors to restrict and monitor access to sensitive clinical data to only those students who had signed required data use agreements. The secondary goals of the computing platform were to support challenges inherent in MOOC education and hosted computing, namely: 1) availability and scalability, 2) secure and privacy preserving, and 3) easy independent access. Given the world-wide access and scope of MOOCs, the platform had to be available across the globe 24 hours a day, 7 days a week, and be able to support potentially hundreds of thousands of learners.^5^ As with all server-based solutions, especially with those hosting clinical data, the platform needed to be secure and have full logging of user activity. Additionally, while it is not legally clear the extent to which MOOCs are subject to the Family Educational Rights and Privacy Act (FERPA)^10^ in order to comply with the course hosting company’s privacy policies, the computing platform needed to preserve student privacy. Finally, given the limited contact with instructors and large student to instructor ratio, the platform onboarding process and use had to be as simple as possible.

### Developing and Deploying the Computing Platform

During development of the computing platform we had a number of resources that shaped the final product. First, we had previously run a version of the specialization as a regular university course that used our university approved Google Cloud platform (GCP)^11^ infrastructure to host student computation. This experience led us to develop a partnership with Google Cloud Healthcare to provide financial support for the creation of the specialization and the hosting of student computation. Second, the Google Cloud Healthcare team routinely hosts healthcare datathons using GCP and MIMIC-III datasets. The hosting guide, system configurations, and other pipelines to support these datathons are all publicly available on GitHub.^12^ Importantly, outside of the choice to use GCP, these resources informed, but did not dictate, the final computing platform created.

GCP is a suite of cloud computing tools that includes support for computing, data storage and databases, networking, identity and security management tools, and advanced analytics for big data applications with all resources organized, managed, and billed to individual projects.^11^ Organizations may have multiple projects and resources can be easily moved between projects as needed. We created a Google Organization (LearnClinicalDataScience) for the computing platform that consisted of two sets of projects - one for developing and prototyping platform improvements (Development) and the other used in production for hosting student work (Production). Within each set of projects, one is devoted to managing student enrollment and platform monitoring (Management) and the other for hosting student work (Computing). All projects use Google Compute Engine^13^ (i.e., virtual servers), Cloud Operations Logging and Monitoring,^14^ and BigQuery.^15^ The Management project handles student enrollment and access with custom R-scripts, Google Forms,^16^ Google Groups,^17^ Cloud Identity and Access Management (IAM),^18^ and SendGrid^19^ email service for student communication. The Computing project hosts student computing using RStudio Server Pro (v1.2.5019-6)^20^ and R (v3.6.3).^21^

We developed a complete version of the computing platform in the Development projects and then performed a series of beta tests. Initial beta tests consisted of project team members (DM, LW) creating user accounts and performing basic computational tasks to ensure that the account management process performed as designed. Basic security checks and penetration tests were conducted by SR to identify any obvious security risks in the platform. We then conducted a group beta test of 27 local users to test simultaneous user registration and to develop an understanding of computing resources required for sample computational workloads similar to those used in the course. Changes to the computing platform and course materials were made following the group beta test and the platform re-tested by project team members (DM, LW). After completion of all beta testing informed improvements, copies of the Development machines were created in the Production projects. The final computing platform was put into production in January 2019. After moving to production, the Development projects were suspended (e.g., shut down, but available for access as needed) and used intermittently to identify the impact of software updates and prototype new platform modifications.

### Evaluating Performance of the Computing Platform

We analyzed data from the first 19 months of computing platform usage (January 15, 2019-August 15, 2020) to understand overall platform performance and associated costs. We identified the number of distinct students who registered for the computing platform (and accepted the data use agreement), and created a frequency map (by country) of all registered students using the city and/or country reported in their signed data use agreement. Computing logs (e.g., unique R sessions, R code inputs, and queries run in BigQuery) were analyzed to determine overall student usage (e.g., number with at least one entry of each type). We also investigated the average volume of platform usage across each day of the week, both with respect to a constant timezone (e.g., overall computing load at any single time point), and student timezone (e.g., what time of day students access course work). All computing logs are captured in UTC. We inferred student local time by mapping student’s reported country/city combination to the regionally observed local time zone. When city level data was not provided, a best attempt to assign a timezone was made by assuming the most prevalent timezone for the area. System performance and reliability were assessed by analyzing all system performance and reliability logs to identify types of errors encountered and total system downtime. Finally, all available discussion forums on the main course hosting site (Coursera), were manually reviewed (DM) and categorized by type of question asked to assess the overall frequency of questions generated by the computing platform. All platform metrics were analyzed with R version 4.0.2 and a variety of packages for data processing, graphing and reporting.^22–25^

## Results

The Clinical Data Science Specialization and the associated course computing platform launched on January 15, 2019. Course programming assignments consist of html-based tutorials with associated RMarkdown documents requiring ∼16MB of storage. As of August 17, 2020 a total of 7,109 students had registered for the first course in the specialization - where students are on boarded to the computing platform. A diagram of the final computing platform implemented is shown in **Figure 1**.

**Figure 1.**
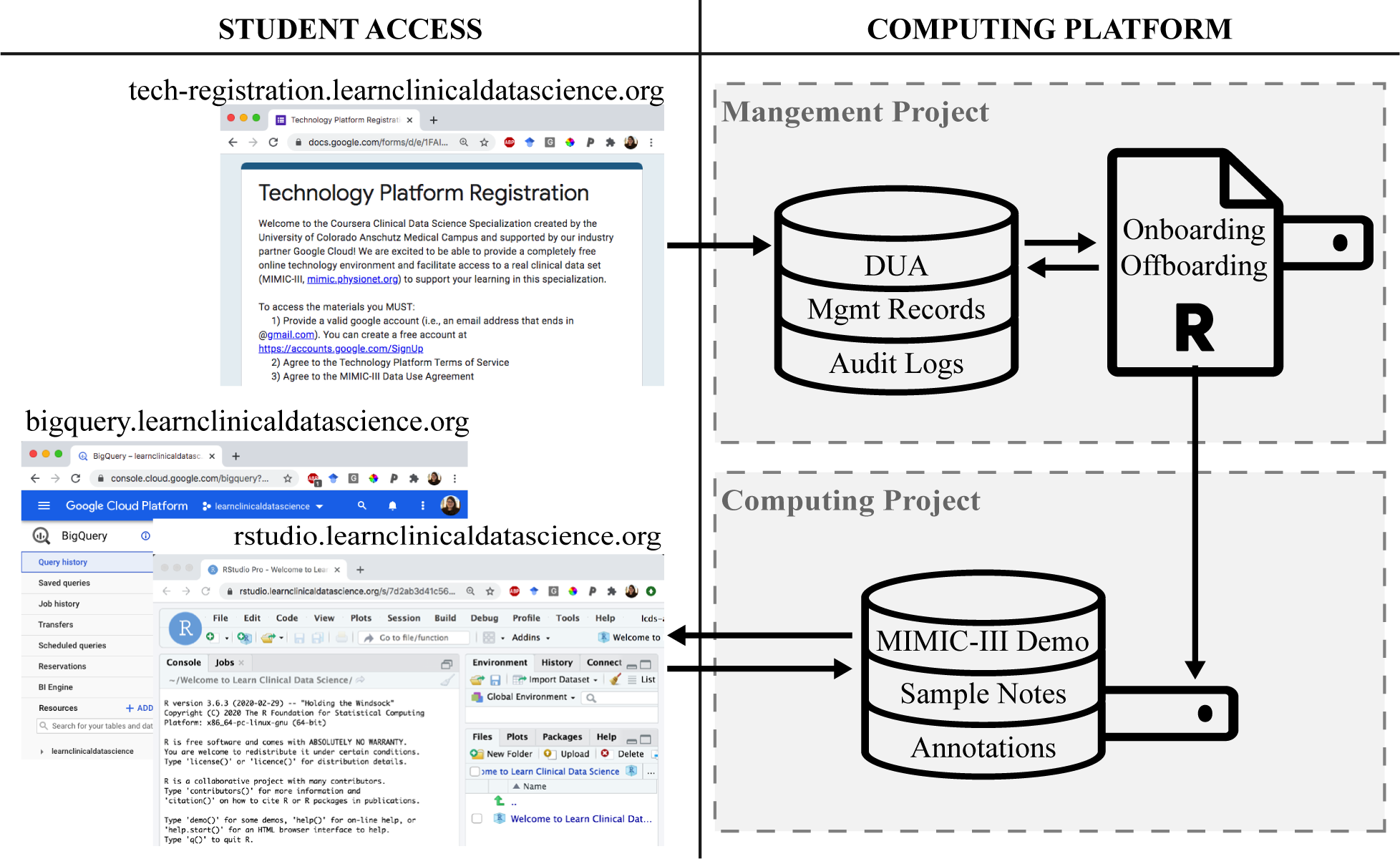
Overview of Computing Platform Design and Student Access Process.

### Computing Platform Technical Details

Each Google project (Management, Computing) consists of one or more virtual servers and a set of associated BigQuery datasets. The Management server has 1vCPU, 3.75GB RAM, and a 50GB SSD boot disk running Ubuntu 18.04. This machine is used to process student enrollments and manage all platform logging activities. The associated BigQuery datasets consist of student management logs (1.2GB), data use agreements (888MB), and R-Session, R-Console, and BigQuery Web UI logs (15.6TB). Earlier versions of the computing platform also used a License Management Server (0.5vCPU, 1.70GB RAM) to host the license key for RStudio Server Pro within the Management Project. All servers in the Management project have firewalls limiting access to the University of Colorado campus. The Computing server has 2vCPU, 7.75GB RAM, a 50GB SSD boot disk and an additional 100GB SSD disk for student file storage. Students complete their coursework on this machine using R and a professional license of RStudio Server. The RStudio Server is configured to have all course related packages pre-installed, restrict terminal access, and set limits on file upload sizes (to attempt to limit loading of non-course data). A firewall is configured such that only https encrypted web traffic on the RStudio interface is accepted. Student’s browsers must support a minimum TLS version of 1.0 and HTTP Strict Transport Security is enabled. Each student has a disk quota (150MB soft, 200MB hard limit). Process time and memory allocation are capped on a per-user basis with calculations limited to 10-min & 2 GB memory. R-sessions are configured with 30 minute idle timeout value. Students have access to 7.9GB of data through BigQuery, including the MIMIC-III demo dataset (in both the original data model and as an OMOP transformation), gold-standard labels for computational phenotyping and NLP courses, and a series of sample clinical notes generated from medical transcription training samples. Every server in the platform has full logging of all system commands and resource usage (i.e., free memory, free CPU cycles, and free disk space) and OS patches are applied automatically at regular intervals.

### Accessing the Computing Platform

Students register for access through a vanity url (tech-registration.learnclinicaldatascience.org) that connects them to a Google Form where they sign the MIMIC-III data use agreement and platform terms of service (e.g., only use the platform for course work, do not attempt to identify other students). These form responses are stored in a Google Sheet that is also accessible from BigQuery. Within the Management Project the Management server runs a custom R-script at 5-minute intervals checking for new registrations. This script determines if the stuwdent is new or returning. New students are assigned a unique student id number, provisioned a linux user account on the Computing server (which is linked to their Google account), added to a private Google Group that has been assigned the appropriate IAM roles for accessing course BigQuery datasets, and a platform expiration date calculated (∼6mo). For returning students (i.e., those with an expired account) their original student id number is identified, their prior linux account reassociated with their Google account, and they are readded to the Google Group. When these onboarding processes are complete, an email is sent to the student confirming their registration and providing links to the course resources. Students can access course data through bigquery.learnclinicaldatascience.org (redirects to the Computing project on BigQuery) and R/RStudio through rstudio.learnclinicaldatascience.org. Students log in to RStudio using their Google account, no passwords or personally identifying information is available on the Computing server. Six months after user registration, the student’s linux account-Google link is removed. While students are not notified of this change, if they try to login they will receive a custom error that asks them to re-register (at this time all their data is preserved).

### Computing Platform Usage

From the launch of the computing platform on January 15, 2019 through August 15, 2020, 2,308 students have requested access to the platform and datasets. Of these registered students, 1,566 (67.9%) have logged in to the computing platform, 1,215 (52.6%) queried course data, and 904 (39.2%) have run R code on the platform. Registered students’ location reported on their data use agreement shows that they live in more than 101 different countries. The overall volume of student registrants by country is provided in **Figure 2**.

Based on data use agreement location, students are accessing the platform from 83 different time zones. With respect to the United States Mountain time zone (MT), the highest total number of connections have occurred at 11pm on Wednesday with 138 connections. The least common hour/day combinations (5pm Saturday, 7pm Friday) have seen only 8 connections total across the 82 weeks the platform has been live. **Figure 3A** shows a summary of the weekly average number of logins by time/day of the week with respect to MT. **Figure 3B** shows a summary of the weekly average number of code commands executed by time/day of the week with respect to the students’ time zone. The most frequent time/ day combination was Thursday at noon, which was driven by work performed by 24 unique students from India (GMT+5:30) across 18 different Thursdays in the evaluation period. Manual review confirmed that this activity was related to course content. Overall, among students who ran R code on the platform, students entered a median of 65.5 [IQR: 11, 187.8] code commands with a single student entering 5,356 code commands. The median amount of storage used by students was < 1MB with a range of 0 - 227.5 MB.

**Figure 2.**
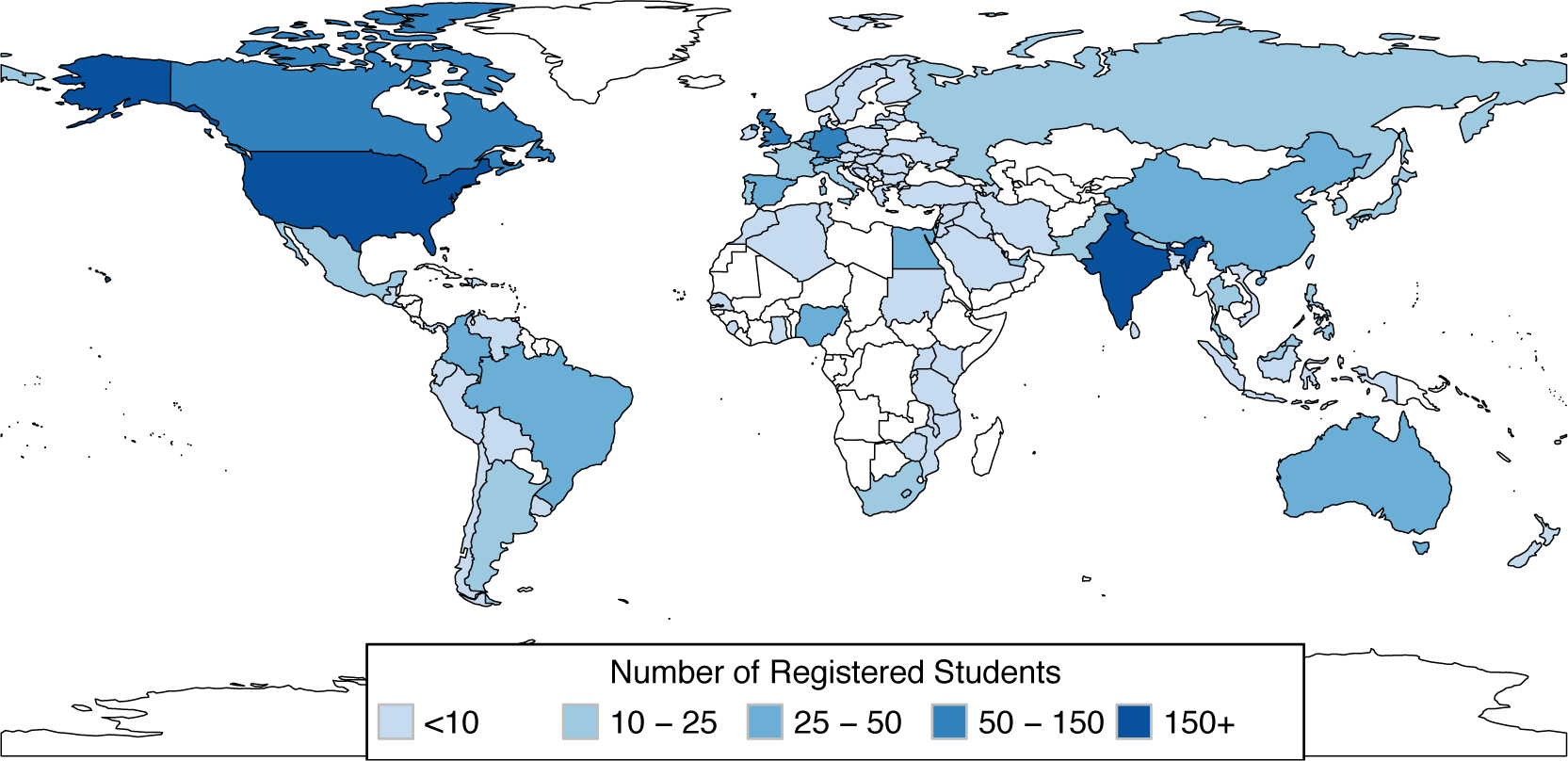
Volume of Computing Platform Registrants by Country.

**Figure 3.**
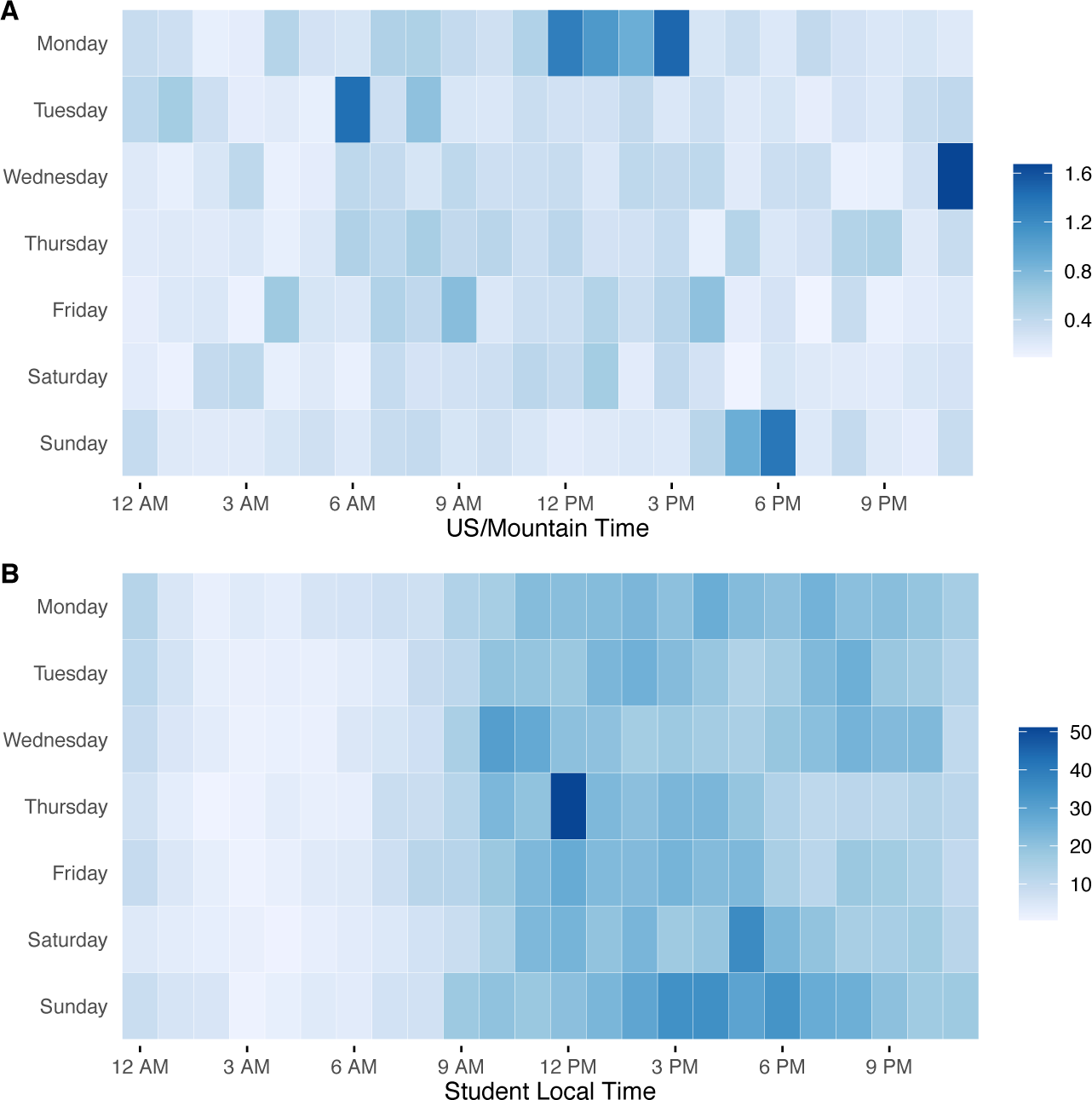
Computing Platform Average Weekly Access. A) This panel shows the weekly average number of R sessions started during each hour/day combination with respectic to the US Mountain Time Zone. B) This panel shows the weekly average number of lines of code entered in the computing platform within each hour/day combination with respect to the student’s local time zone.

### Computing Platform Performance

Over the 82 weeks the computing platform has been running there were 25 unexpected alerts or other errors, of which 6 resulted in downtime where students were unable to access the system. The majority of alerts originated on the Computing server (n=18), 13 alerts were related to CPU and memory utilization (i.e., available CPU < 50%, free RAM < 25%). Five uptime alerts on the Computing server were related to: DNS outages (n=2, downtime only for affected regions), RStudio Server errors (n=2), and a License Management Server outage (n=1). The License Management Server had 5 unexpected outages totaling just under 3 hours of downtime, however the majority (n=4) of these outages lasted less than 30min - the interval the Computing server uses to confirm an active licence. One incident occurred overnight in MT and accounted for approximately an hour and a half of downtime on the Computing server and resulted in at least two students unable to access the machine. The Management server had no unscheduled downtime. **Table 1** provides a summary of the total number of unexpected alerts and computing platform outage durations. Overall the computing platform has had a system uptime percentage of 99.8%.

**Table 1.** Summary of Computing Platform Alerts and Outages.

|  | Count<br>Unexpected<br>Alerts | Alert/Outage<br>Duration | Computing<br>Platform Outage<br>Duration |
| --- | --- | --- | --- |
| License Management Server | 5 | 2hr 54min | 1hr 24min |
| Computing Server |  |  |  |
| CPU Utilization Alert | 11 | 5hr 33min | - |
| Memory Utilization Alert | 2 | 183hr 16min | - |
| Uptime Alert | 5 | 23hr 10min | 22hr 7min |
| *Total Computing Platform Unavailability:* |  |  | 23hr 31min |

There have been 13 instances of user onboarding errors. First, although we require all students to register with an “@gmail.com” address, some accounts are treated in IAM as an “@googlemail.com” address. One student registered with such an account and was unable to access the platform until we adjusted our onboarding process to account for this scenario. Three students registered with the same Google account multiple times within 5 minutes (the onboarding script interval) which resulted in them unexpectedly being assigned to the same user account. When multiple registrations occur in the same batch, a student id was only assigned to the first registration with subsequent registrations labelled as NA, however an account and account/mapping were performed for both accounts. As the RStudio-Google authentication map uses the most recently added mapping, all students accessed a single (shared) “NA” user account. Finally, 9 students were not initially granted data access when registering. This was due to exceeding a Google limit on IAM users that can share a single role (n=1,500).

Across the four launched courses of the Clinical Data Science Specialization there were 196 forum posts available for analysis, with 259 questions/issues raised (students can comment on forum posts to answer a question or echo concerns). Of these issues, 61 (23.5%) related to the computing platform with the majority (n=38, 62.3%) due to the students not registering for access. The remaining 23 included technical issues with students’ registration (n=10) or students who had issues locating the resources they needed (n=13).

## Discussion

Managing student computing environments is a major challenge in data science education which is magnified when dealing with MOOCs and sensitive clinical data. When developing a series of clinical data science MOOCs we identified the need to develop a computing platform with the primary objective of restricting access to and logging access of sensitive health data. Here we present the results of that work - a computing platform that can only be accessed by students who have completed all requirements for data access (signed data use agreement) where all data access (e.g., SQL queries) and analysis (e.g., R-console commands) are logged and available for review. In addition to these fundamental objectives, we also had three technical requirements related to perceived challenges with hosted computing in MOOCs: 1) availability and scalability, 2) secure and privacy preserving, and 3) easy independent access.

Although we prepared our courses with the potential for an international audience, it wasn’t well known whether clinical data science coursework with a particular focus on customs/regulations in the United States would be of global interest. However given the global popularity of other data science MOOCs^5^, we designed the platform to be accessible worldwide. Indeed, we had students from around the globe register for access and since deployment of the platform at least one student has started an R session at every single hour of every day of the week. The global use of the platform is confirmed by analyzing platform usage relative to the students’ time zone, where most students access the platform during more traditional learning hours (9am-midnight). This platform has also proven to be very stable with only 6 incidents resulting in system-wide outages for only 23.5 hours across the 13,896 hours the platform has been available.

One of the benefits of using a cloud infrastructure is that the platform has proven to be scalable on demand. Initial deployments used larger servers and directory storage (500GB), as actual platform usage started we were able to initially reduce our storage to 100GB and eventually increase it to 150GB after a year of additional student access. Similarly, we can adjust the server specifications (CPUs and RAM) as needed with limited downtime (simply requires restarting the server). This flexibility has had the additional benefit of allowing for cost control measures as we can shrink and grow the machine as needed. An additional benefit has been the query caching function within BigQuery where identical repeated queries are not charged. These types of repeated queries are common for students working through set exercises. Even without aggressive system size optimization, the computing platform development cost $1,431 and continuous access for more than 2300 students across 19 months on the production platform only cost $2,941.

We took multiple steps to ensure the system security and student privacy. First, our platform design uses standard web-based security steps including enabling SSL and limiting server access for students as much as possible beyond those resources required for completing course content. In addition to logging all access and activity, we routinely monitor those logs for unapproved access attempts including running system commands. Even if a student somehow bypassed our permissions restricting access for viewing/modifying system files, all student files are labeled by student number only. Additionally, although we use Google Groups for managing data access roles, by using an Organization Google account the membership of this group (and group enrollment status) is kept entirely hidden from students. To our knowledge, outside of the single instance of three students getting mapped to the same user account, no private student information has been inappropriately accessed.

Given the exceptionally high student to instructor ratios in MOOCs, it was critical that our platform be accessible without extensive instruction or hands-on attention. To this end students interact with only three sites, all with custom vanity URLs - one for registration and two for platform access. By all available measures our approach has worked well. Technical issues accounted for a minority of forum posts across the courses. The majority of the issues raised were related to students having not attempted to register - suggesting that the primary improvement is needed within our learning management system to highlight the need to access the external site. Although a minority of students did have actual technical issues accessing the site due to registration errors, these issues were usually fixed within the same or subsequent day after reporting on the forum. Although available data supports the easy accessibility of the platform, the number of students registered is dramatically lower than course registrants. It is possible that this drop is due to unreported student issues with the platform. Alternatively, MOOCs experience a well known phenomena where the number of students registering far exceeds those who perform any course work with even fewer completing the course.^26^

While we have pleasantly been surprised by the overall efficacy and efficiency of the system, there are numerous limitations to our approach. First, and most importantly, this approach was not without cost. Although as highlighted above the expenses are manageable by right-sizing the system size, there is a non-trivial cost associated with hosting student computation. Our numbers are also much smaller than would be expected for data science courses that use much larger datasets/computationally intensive algorithms. The MIMIC-III demo data contains the records of only 100 patients and though the sample note corpus is larger, most of the computation performed on the platform is happening within the database. Second, although there is renewed interest in online computing solutions due to the COVID-19 pandemic requiring remote education, this solution is likely over-engineered for the majority of university courses. Additionally, due to platform security concerns we have not made the processing scripts available publically.

Finally, we have found a number of unexpected benefits from the course platform. First, logistically having full logging of all issues commands has been invaluable when answering student questions, both technical and around course content. When students report that they can’t access course resources it’s easy to pinpoint whether they simply haven’t registered or if they are having some other issue. For course problems we are able to see what commands they are running and provide targeted recommendations for “common issues” students encounter. Second, as educators and content creators, these logs also allow for valuable insight into how students interact with course materials. We hope to perform more robust studies of these data in the future to inform both clinical data science education best practices and to improve our own course materials.

## Conclusion

We have created a computing platform to support clinical data science MOOC education that has been scalable, globally available, secure, privacy preserving, and generally supported independent access by a large number of students.

## Acknowledgements & Funding

We thank our Google partners, especially Marianne Slight, Kate Strasburger, and Stuart O’Brian for their support and their team’s technical advice. We are especially grateful to the MIT Laboratory for Computational Physiology whose work and support allowed us to provide students with real clinical data. Our beta testers and students who have provided feedback on the platform have all significantly improved our final production platform. Finally we would like to thank the extraordinary team who have contributed to the creation of the clinical data science MOOC: Chan Voong, Christine Mousavi, Jay Billups, Janet Corral, Deborah Keyek-Franssen, Jill Taylor, Jill Lester, Jaimie Henthorn, Aileen Sanders, Alesia Blanchard, Ashley Boshoven, and Benita Bazemore-Cook. Computing platform use and development costs were supported by our partnership with Google Cloud Healthcare, and complimentary RStudio Server Pro licenses were provided by the RStudio Education team.

## References

1. Kross S, Guo PJ. Practitioners Teaching Data Science in Industry and Academia: Expectations, Workflows, and Challenges. 2019 May 2 [cited 2020 Aug 18];1–14. Available from: https://dl.acm.org/doi/pdf/10.1145/3290605.3300493

2. RStudio Cloud - Do, share, teach, and learn data science [Internet]. [cited 2020 Aug 19]. Available from: https://rstudio.cloud/

3. Suen A, Norén L, Liang A, Tu A. Equity, Scalability, and Sustainability of Data Science Infrastructure. In: Proceedings of the 17th Python in Science Conference doi [Internet]. 2018. Available from: http://conference.scipy.org/proceedings/scipy2018/pdfs/anthony_suen_laura_noren_alan_liang_andrea_tu.pdf

4. Leek J. The Data Scientist’s Toolbox [Internet]. Coursera. Available from: https://www.coursera.org/learn/data-scientists-tools?specialization=jhu-data-science

5. Kross S, Peng RD, Caffo BS, Gooding I, Leek JT. The Democratization of Data Science Education. Am Stat [Internet]. 2020 Jan 2;74(1):1–7. Available from: https://doi.org/10.1080/00031305.2019.1668849

6. Brooks C. Introduction to Data Science in Python [Internet]. Coursera. Available from: https://www.coursera.org/learn/python-data-analysis?specialization=data-science-python

7. Bresnick J. Lack of Talent, Direction Afflict Healthcare Data Analytics Plans [Internet]. Health IT Analytics. [cited 2020 Aug 19]. Available from: https://healthitanalytics.com/news/lack-of-talent-direction-afflict-healthcare-data-analytics-plans

8. Learn Clinical Data Science [Internet]. [cited 2020 Aug 19]. Available from: https://www.learnclinicaldatascience.org/

9. Johnson A, Pollard T, Mark R. MIMIC-III Clinical Database Demo [Internet]. PhysioNet; 2019. Available from: http://dx.doi.org/10.13026/C2HM2Q

10. Young EM. Educational Privacy in the Online Classroom: FERPA, MOOCS, and the Big Data Conundrum. Harvard Journal of Law and Technology [Internet]. 2015 [cited 2020 Aug 21]; Available from: https://www.semanticscholar.org/paper/5bcd4885d7cd291fcae72c05c802338869d55859

11. Google Cloud Computing, Hosting Services & APIs [Internet]. [cited 2020 Aug 21]. Available from: https://cloud.google.com/gcp/

12. Datathon Support by Google Cloud Healthcare [Internet]. GitHub. 2019 [cited 2020 Aug 21]. Available from: https://github.com/GoogleCloudPlatform/healthcare

13. Compute Engine: Virtual Machines (VMs) [Internet]. [cited 2020 Aug 21]. Available from: https://cloud.google.com/compute

14. Operations: Cloud Monitoring & Logging [Internet]. [cited 2020 Aug 21]. Available from: https://cloud.google.com/products/operations

15. BigQuery: Cloud Data Warehouse [Internet]. [cited 2020 Aug 21]. Available from: https://cloud.google.com/bigquery

16. Google Forms: Free Online Surveys for Personal Use [Internet]. [cited 2020 Aug 21]. Available from: https://www.google.com/forms/about/

17. Google Groups [Internet]. [cited 2020 Aug 21]. Available from: https://groups.google.com/

18. Cloud Identity and Access Management [Internet]. [cited 2020 Aug 21]. Available from: https://cloud.google.com/iam

19. Email Delivery Service [Internet]. [cited 2020 Aug 21]. Available from: https://sendgrid.com/

20. RStudio Server Pro [Internet]. [cited 2020 Aug 21]. Available from: <https://rstudio.com/products/rstudio-server-pro/>

21. R Core Team. R: A Language and Environment for Statistical Computing [Internet]. Vienna, Austria: R Foundation for Statistical Computing; 2013. Available from: http://www.R-project.org/

22. Wickham H, Averick M, Bryan J, Chang W, McGowan L, François R, et al. Welcome to the Tidyverse. JOSS [Internet]. 2019 Nov 21;4(43):1686. Available from: https://joss.theoj.org/papers/10.21105/joss.01686

23. Neuwirth E. RColorBrewer: ColorBrewer Palettes [Internet]. 2014. Available from: https://CRAN.R-project.org/package=RColorBrewer

24. Kassambara A. ggpubr: “ggplot2” Based Publication Ready Plots [Internet]. 2020. Available from: https://CRAN.R-project.org/package=ggpubr

25. South A. rworldmap: A New R package for Mapping Global Data [Internet]. Vol. 3, The R Journal. 2011. p. 35–43. Available from: http://journal.r-project.org/archive/2011-1/RJournal_2011-1_South.pdf

26. Reich J, Ruipérez-Valiente JA. The MOOC pivot. Science [Internet]. 2019 Jan 11;363(6423):130–1. Available from: http://dx.doi.org/10.1126/science.aav7958

